# The influence of soft tissue volume on estimates of skeletal pneumaticity: implications for fossil archosaurs

**DOI:** 10.1101/2024.10.08.617080

**Authors:** Maria Grace Burton, Juan Benito, Kirsty Mellor, Emily Smith, Elizabeth Martin-Silverstone, Patrick O’Connor, Daniel J. Field

## Abstract

Air space proportion (ASP), the volume fraction in bone occupied by air, is frequently applied as a measure for quantifying the extent of skeletal pneumaticity in extant and fossil archosaurs. Nonetheless, ASP estimates rely on a key assumption: that the soft tissue mass within pneumatic bones is negligible, an assumption that has rarely been explicitly acknowledged or tested. Here, we provide the first comparisons between estimated air space proportion (where the internal cavity of a pneumatic bone is assumed to be completely air-filled) and true air space proportion (ASPt, where soft tissues present within the internal cavities of fresh specimens are considered). Using birds as model archosaurs exhibiting postcranial skeletal pneumaticity, we find that estimates of ASPt are significantly lower than estimates of ASP, raising an important consideration that should be acknowledged in investigations of the evolution of skeletal pneumaticity and bulk skeletal density in extinct archosaurs, as well as in volume-based estimates of archosaur body mass. We advocate for the difference between ASP and ASPt to be explicitly acknowledged in studies seeking to quantify the extent of skeletal pneumaticity in extinct archosaurs, to avoid the risk of systematically overestimating the volume fraction of pneumatic bones composed of air.

## Introduction

Skeletal pneumaticity refers to the presence of epithelial-lined, air-filled cavities within bones. The presence of ‘hollow bones’ in many birds is among the most distinctive aspects of extant avian biology, and has been a subject of interest for centuries [e.g., 1, 2–4]. Birds are the only extant tetrapods to exhibit postcranial skeletal pneumaticity (PSP), although osteological correlates of PSP (e.g., pneumatic foramina through which diverticula of the respiratory system invade the skeleton) are observed in several groups of extinct ornithodiran (bird-line) archosaurs. These groups (pterosaurs, sauropodomorph dinosaurs, and non-avian theropod dinosaurs including some Mesozoic avialans) document a deep evolutionary history of PSP, dating back to at least the Late Triassic (∼210 Ma) [e.g., 5, 6–16]. Recent evidence suggests that PSP may have evolved at least three times independently among those groups [17]; however, the presence of an air-sac system exhibiting invasive diverticula is thought to be homologous among these groups, and generally similar to that observed in extant birds [e.g., 7, 10, 11, 15, 18–23].

Variation in the presence and extent of skeletal pneumaticity is thought to have facilitated the adaptive decoupling of standard mass-volume relationships throughout the evolutionary history of Ornithodira [24]. Understanding the biological importance of PSP in different groups is dependent on a clear understanding of differences in the extent of pneumaticity among different taxa; nonetheless, such inferences in long-extinct groups are challenging. As the only extant group to exhibit PSP, birds have generally provided a model for investigations of the pneumatic systems of extinct non-avian archosaurs [e.g., 11, 12, 14, 18], and patterns among birds have provided insight into the evolutionary and developmental origins of pneumaticity [e.g., 10, 17, 24, 25], as well as function of the pneumatic system [e.g., 9, 24, 26, 27–31].

Air Space Proportion (ASP) is a measure of pneumaticity proposed by Wedel [8], defined as “the proportion of the volume of a bone—or the area of a bone section—that is occupied by air spaces.” Since its inception, a recognised limitation of ASP is the assumption of negligible soft tissue volume within the internal cavities of pneumatic bones (i.e. the assumption that the entire internal cavity of a bone is completely air-filled). Nonetheless, a recent investigation of humeral pneumaticity across a wide phylogenetic sample of extant birds revealed variable amounts of soft tissue within the internal cavities of pneumatic extant bird humeri: 26% of the volume fraction within the internal cavity of pneumatic humeri was occupied by intraosseous soft tissues (e.g., adipose tissue) on average, rather than 0% as would be assumed by the uncritical application of ASP [30]. Indeed, the volume fraction of intraosseous soft tissues within a pneumatic humerus was as high as 71% in that study, in a specimen of the Common Swift (*Apus apus*). These findings suggest that the extent of pneumaticity as estimated by ASP has been systematically overestimated in previous investigations, yet it remains unclear whether incorporating more realistic values of soft tissue within the internal cavities of pneumatic fossil bones would meaningfully influence estimates of the extent of skeletal pneumaticity, bulk bone density, and body mass in fossil archosaurs.

In this study, we build on the dataset of Burton et al. [30], using fresh specimens of extant birds with intact soft tissues to compare measures of the extent of pneumaticity excluding soft tissues (ASP) with an equivalent measure of ‘true’ air space proportion that includes soft tissues (ASPt) in select pneumatic bones. We use previously published data on humeral pneumaticity [30] for the bulk of our analyses, and include other skeletal elements (e.g., cervical and thoracic vertebrae, femora) as case studies for comparison with previously published work. We aim to clarify whether the assumption of negligible soft tissue within the internal cavities of pneumatic bones is justifiable, and if not, address the implications this may have for previous findings from the fossil record (e.g., mass estimation in fossil taxa with pneumatic elements; Wedel [8]). We also aim to understand the feasibility of estimating ASP and ASPt from fossil or dry skeletal material, including addressing estimates gathered from fragmentary sections of long bones.

## Methods

### (a) Taxon sampling

We sampled a total of 41 extant bird species with representatives from 17 ordinal-level clades. Of that sample, our humeral analyses reinvestigated the sample of 36 species from 15 ordinal-level clades from Burton et al. (2023). Sampled specimens were sourced from the University of Cambridge Museum of Zoology (UMZC), except for *Eudromia elegans*, which was provided by the Royal Veterinary College (RVC). Specimens consisted of deceased, frozen, externally intact, mature individuals of which the vast majority were salvaged. Sex of specimens was not a controlled factor, and in most cases were unknown. Due to the nature of working with salvaged specimens, time between death and freezing were uncontrolled factors. Potential effects are discussed below. See supplementary table S1 for complete data on specimens sampled.

### (b) Data collection

Complete frozen bird specimens were microCT scanned with a Nikon XTEK H 225 ST scanner at the Cambridge Biotomography Centre (CBC), and all scanned material was digitally segmented and rendered three-dimensionally using VGSTUDIO MAX 3.4.5 or 2023.2.1 (Volume Graphics, Heidelberg, Germany). Pneumatic elements were identified in CT scans by clear invasion of air (which appears black in CT images) into the internal cavity of a bone. We followed the methodology of Burton et al. [30] to generate 3D volumetric reconstructions of the regions of pneumatic elements occupied by bone, soft tissues, and air (Fig. 1).

**Figure 1.**
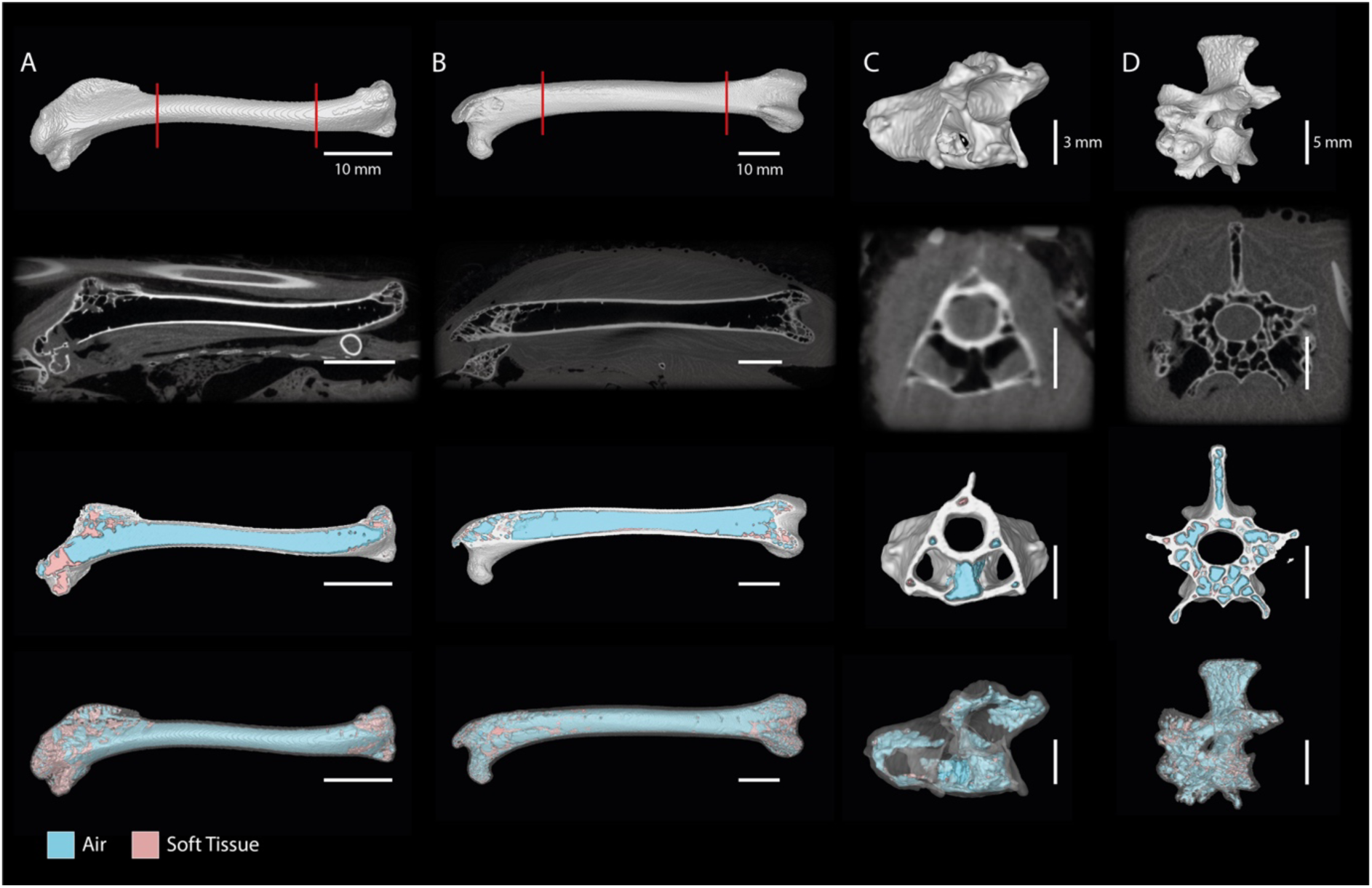
Computed Tomography (CT) slice data (second row) and digital polygon reconstructions of elements investigated in this study. Top row, 3D reconstructions of bones. Second row, representative CT slice through element. Third row, 3D reconstruction of element sliced at point shown in the second row, allowing for visualisation of segmented air (blue) and soft tissues (pink). Fourth row, 3D reconstruction of whole element showing segmented air (blue), soft tissues (pink), with the bone rendered semi-transparent. (A) right humerus of *Pitta moluccensis*; (B) left femur of *Phasianus colchicus*; (C) fifth cervical vertebra of *Anas crecca*; (D) second thoracic vertebra of *Phasianus colchicus*. The region between the red lines in the top row represents the section of the humeral and femoral diaphyses examined in our diaphysis vs. whole bone comparison (supplementary table S1). Scale bars represent 10 mm (A and B), 3 mm (C) and 5 mm (D).

We initially segmented any bone material comprising the element of interest using manual segmentation bound by grey value thresholds that were locally appropriate. Using a closing operation on our ‘bone’ model and some manual segmentation to fill any remaining holes, we created a ‘total volume’ model from which air within the internal cavity was segmented using grey value thresholds appropriate for the scan. We subsequently segmented any residual soft tissues (collectively labelled ‘marrow’) that did not clearly fall within the density ranges of ‘bone’ or ‘air’ within the ‘total volume’ model. Grey scale ranges were not standardised across scans, thus appropriate thresholds for ‘bone’ and ‘air’ were decided on a scan-by-scan basis based on visual assessment of their observed boundaries in CT scans (see supplementary Table S1 for all grey value ranges used to create models). For ‘total volume’ models of cervical vertebrae, we exclude the enclosed space of the transverse foramina, which is bounded laterally by the costotransverse handle (ansa costotransversaria; [32]), yet open on their cranial and caudal ends, as this space is not truly intraosseous and therefore not considered a site of skeletal pneumatisation. For ‘total volume’ models of both cervical and thoracic vertebrae, we also exclude the neural canal space for the same reason. Due to some areas of the ‘marrow’ region of interest (ROI) likely representing transition voxels in the CT scan between bone (white) and air (black), we followed the method of Burton et al. [30] by performing an erode/dilate (i.e. opening) operation on the originally segmented ‘marrow’ ROIs. We slightly modify this method by taking the volume lost through the opening operation on the original ‘marrow’ ROI and distributing that lost volume equally between ‘bone’ and ‘air’ to give their final corrected volumes. In this way, the original total volume of the element is unchanged, and we reduced the chance of overestimating the true amounts of soft tissue included in the ‘marrow’ volume, as we only included non-negligible voxel groups in our corrected ‘marrow’ component. Preliminary tests of this additional methodological step show that mean ASPi (air space proportion of the internal cavity [V_air_ / V_air + marrow_]; [30]) and ASPt of pneumatic humeri data from Burton et al. [30] remained virtually unchanged (mean ASPi = 0.75 with the new method as opposed to 0.74; mean ASPt = 0.43 with both methods), and therefore results and discussions from that study likely remain consistent with the results from the present work.

Using our volumetric data we calculated ASP as originally described—assuming the internal cavity of pneumatic elements to be completely air-filled—by combining air and marrow volumes to give the internal cavity volume and dividing that by the total volume of the element (i.e. V_air + marrow_ / V_air + marrow + bone_). We also calculated ASPt, as the volume of the air component divided by the total volume of the element (i.e. V_air_ / V_air + marrow + bone_; see Table 1 for air space proportion definitions used in this study).

**Table 1.** Summary of air space proportion acronyms and definitions as used in the context of this study. ‘V’ represents ‘volume’. Note: ‘V_marrow_’ is a generalisation for any intraosseous soft tissue volume, not specific to marrow.

| Acronym | Definition | Calculation |
| --- | --- | --- |
| ASP (air space proportion) | The proportion of the volume of a bone that is occupied by air spaces, assuming the entire internal cavity of a pneumatic bone is air-filled. | $\frac{V_{air} + V_{marrow}}{V_{bone} + V_{air} + V_{marrow}}$ |
| ASPt (true air space proportion) | The proportion of the volume of a bone that is occupied by air spaces. | $\frac{V_{air}}{V_{bone} + V_{air} + V_{marrow}}$ |
| ASPi (air space proportion of the internal cavity) | The proportion of the volume of the internal cavity of a pneumatic bone that is occupied by air spaces. | $\frac{V_{air}}{V_{air} + V_{marrow}}$ |

Most volumetric data from pneumatic humeri were taken from Burton et al. [30], although we also collected similar data on only the diaphysis (shaft) of each humerus to enable comparisons between ASP and ASPt to reflect a situation in which only the diaphysis was available (e.g., similar to that in fragmentary fossil long bones). We added data on pneumatic avian femora, a mix of anterior, mid- and posterior cervical vertebrae and anterior thoracic vertebrae. These additional elements were sampled as case studies to provide pilot comparative data to our considerably larger humeral sample. See supplementary table S1 for full details on elements, volumes, ASP and ASPt variables for each specimen.

Limitations to this method of collecting volumetric air, marrow and bone volumes using CT data derived from frozen specimens are the same as discussed in Burton et al. [30] and their supplementary information, though we note that the effects of decomposition on pneumatic spaces appear to affect vertebrae at a faster rate than the humeri in the same sample. As such, several specimens that were included in our humeral sample were deemed unsuitable for reliable data collection on pneumatic vertebrae.

We also estimated the true bulk density of each sampled element with soft tissues included (i.e. as in the ASPt measure) and excluded (i.e. as is assumed in the ASP measure). Bulk density has also been referred to as ‘specific density’ [e.g., 33], and when given in g cm^-3^ is numerically equivalent to estimates of ‘specific gravity’ (SG) in some previous literature [e.g., 8, 18, 34]. To calculate bulk density from our raw measurements of bone, marrow and air volume, we follow the method described by Burton et al. [30], where the estimated density of avian bone was assumed to be 2.05 g cm^-3^ [35], that of marrow was assumed to be the same as water at 1 g cm^-3^ [30, 36], and that of air was assumed to be negligible. For the bulk density estimate using ASP, marrow and air were treated equivalently (0 g cm^-3^).

### (c) Statistical analysis

A paired samples *t*-test was used to assess whether a statistically significant difference exists between the ASP and ASPt measures obtained from avian humeri. Two additional paired samples *t*-tests were carried out to compare whether there are statistically significant differences for each measure between the humerus as a complete whole compared to just the diaphysis (shaft). Where significant differences were identified from *t*-tests, we also calculated Cohen’s *d* [37] for paired samples as a measure of effect size. Spearman’s rank correlation was used to assess the relationship between ASP and ASPt among complete humeri. All statistical tests were carried out in *R* [38], with *R* ‘stats’ package used for *t*-tests and Spearman rank analyses, and the ‘effsize’ package [39] for calculating Cohen’s *d*.

For the remaining elements (femur, cervical and thoracic vertebrae) included here as case studies, statistical analyses were not conducted due to limited sample size. However, mean values were compared at face value to give some indication of the discrepancy between ASPt and ASP.

## Results

### (a) Humeral ASP versus ASPt

Among complete humeri, ASP ranged between 0.33 in *Ensifera ensifera* to 0.74 in *Chauna torquata* with a mean of 0.57. ASPt ranged between 0.13 in *Apus apus* to 0.65 in *Asio flammeus* with a mean of 0.43 (Table 2). Among paired differences between ASP and ASPt, the smallest observed difference was 0.01 (*Anas crecca*) and the greatest was 0.28 (*Perdix perdix*). We detected a significant difference in air space proportion when excluding and including soft tissue proportions (paired *t*-test: *t*(37) = 11.594, p = 7.017e-14), with the mean of the paired differences being 0.14 (CI 95%: 0.11 to 0.16). This indicates that when soft tissues are included in the ASP measure, the estimated air space proportion is reduced by a factor of 0.14 on average. In addition, Cohen’s effect size value (*d* = 1.24) indicates a large effect (*d* > 0.8; [37]).

**Table 2.**
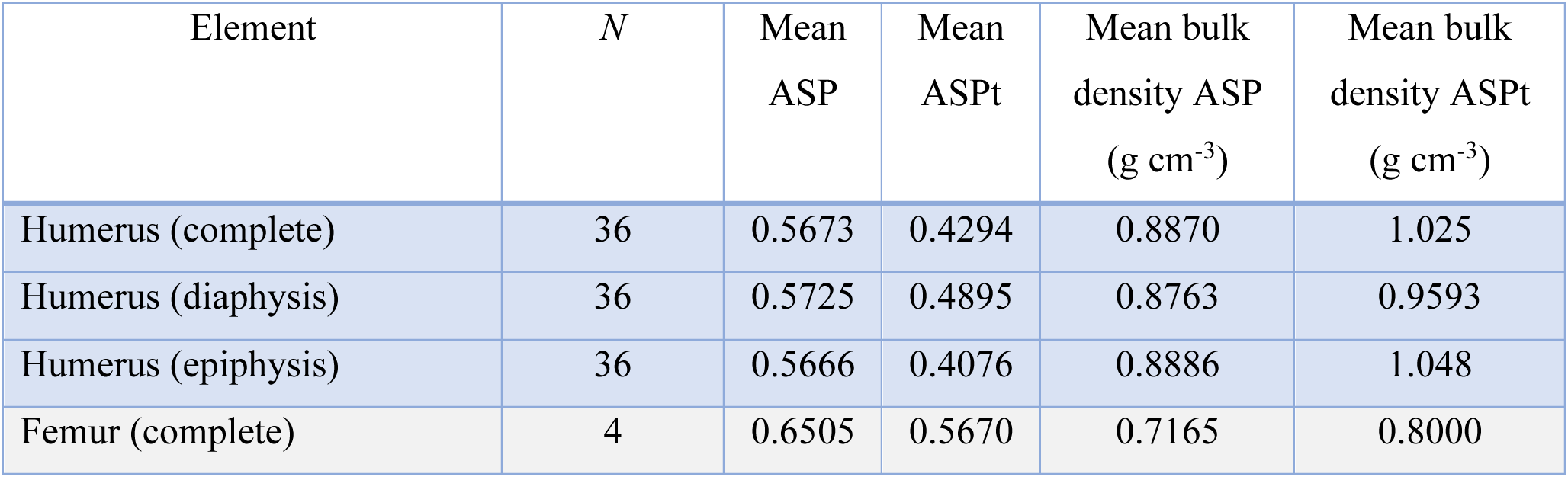
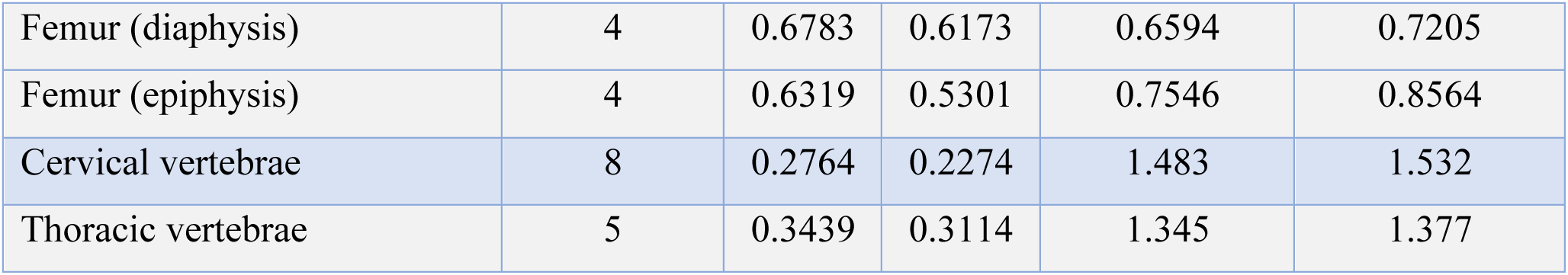
Mean ASP and ASPt calculations by element, as well as mean bulk density estimates with soft tissues excluded (ASP) or included (ASPt). Mean values are rounded to four significant figures.

Differences in bulk density estimates according to the exclusion versus inclusion of soft tissues (i.e. using ASP versus ASPt, respectively) are inversely proportional to the differences between the ASP and ASPt measures of the complete element themselves, due to the assumption of marrow density being equivalent to water (1 g cm^-3^). Therefore, paired samples *t*-tests between the humeral bulk density estimates provide redundant information here. However, effect size may still vary despite identical *t*-test results, and the effect size for the differences in humeral bulk densities according to ASP and ASPt indicates a medium effect (Cohen’s *d* = 0.70; [37]). Together, these results indicate that excluding soft tissues significantly increases estimates of air space proportion in pneumatic bones, and the practical significance of this exclusion is that of a medium to large effect.

A Spearman’s rank correlation was conducted to evaluate the relationship between ASP and ASPt among pneumatic humeri. The coefficient indicated a strong, positive and statistically significant correlation between the two variables (ρ(36) = 0.758, p = 3.314e-07). We can interpret this to mean that generally, as ASP increases, ASPt also increases such that when ASP is relatively high, ASPt also tends to be relatively high (and *vice versa* for relatively low proportions). However, their rankings within our dataset are not always identical due to the variable proportions of soft tissues within the internal cavity of the pneumatic elements. Where large discrepancies between the ASP and ASPt rankings within the dataset are present, it may indicate the presence of more or less soft tissues than expected, depending on the direction of the change in rankings. For instance, taxa with humeri whose rankings among ASPt were ten (∼25% of the dataset) or more places lower than their rankings for ASP are the partridge *Perdix perdix* (18 ranks), the owl *Strix aluco* (17 ranks) and the sparrowhawk *Accipiter nisus* (15 ranks). For these cases, it would imply that these taxa exhibit less air, or more soft tissues, within their humeri than expected relative to the rest of the dataset. On the other hand, taxa representing humeri whose rankings among ASPt were ten or more places higher than their rankings for ASP are the skua *Stercorarius antarcticus* (15 ranks), the parrot *Lorius garrulus* (11 ranks) and the partridge *Alectoris rufa* (10 ranks). In these cases it would imply that these taxa exhibit more air, or less soft tissues, than expected relative to the rest of the dataset.

### (b) Preliminary results from additional skeletal elements

We collected 3D volumetric data from five pneumatic femora representing four extant bird species. Among complete femora, ASP ranged from 0.55 in the pheasant *Phasianus colchicus* to 0.76 in the flamingo *Phoenicopterus roseus* with a mean of 0.65. ASPt of complete femora ranged from 0.45 in one femur of the grouse *Lagopus lagopus scotica* to 0.76 in *P. roseus* with a mean of 0.57 (Fig. 2). However, such mean values should be interpreted with caution due to the small sample size on which they are based, a result of the fact that a far greater proportion of extant bird species exhibit pneumatic humeri than pneumatic femora. More importantly here may be the paired differences between ASP and ASPt. The greatest paired difference between ASP and ASPt of complete femora was 0.18 in *L. lagopus scotica*, with the second femur from this individual exhibiting a relatively high difference of 0.14 as well. The remaining femora exhibited paired ASP to ASPt differences of 0.00 to 0.05, with the lowest of those paired differences (0.00) in *P. roseus*. Among all the elements examined in our entire dataset, the femur of *P. roseus* exhibits the greatest degree of pneumaticity, where ASP matches ASPt due to a complete lack of detectable soft tissue within its internal cavity. The mean of the paired ASP to ASPt differences among complete femora was 0.08. Despite our small sample size precluding a statistical assessment (e.g., Spearman’s rank correlation) of the relationship between ASP and ASPt in femora, there seems to be a general trend of a direct relationship between ASP and ASPt.

**Figure 2.**
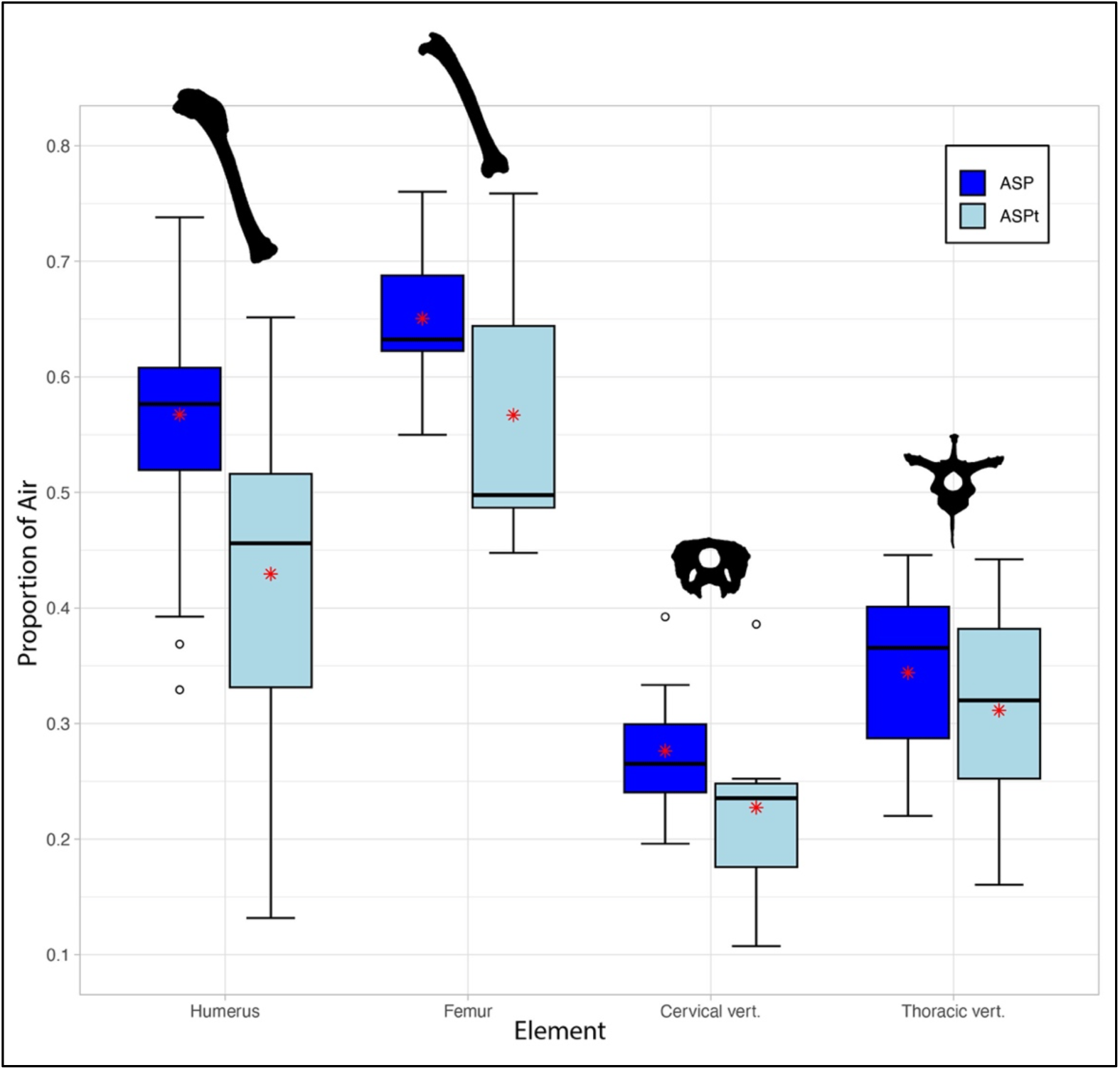
Box and whisker plot showing ASP (dark blue) versus ASPt (light blue) for representative elements included in this study. Bold lines on box plots represent median values. Upper whiskers correspond to data within 1.5 times the interquartile range over the 75th percentile, lower whiskers correspond to data within 1.5 times the interquartile range under the 25th percentile. Red star represents the mean. Humerus silhouette: *Pitta moluccensis*; femur silhouette: *Phasianus colchicus*; cervical and thoracic vertebrae silhouettes: *Callonetta leucophrys* (not to scale).

We collected similar data on pneumatic cervical and thoracic vertebrae. For cervical vertebrae (*n* = 9 from eight different species), ASP ranged from 0.20 in an anterior cervical of the pygmy goose *Nettapus auritus* to 0.39 in a posterior cervical of the rook *Corvus frugilegus*, with a mean ASP value of 0.28 (Fig. 2). For ASPt of the same cervical vertebrae sample, ASPt ranged from 0.11 in the same anterior cervical of *N. auritus* to 0.39 in the same posterior cervical of *C. frugilegus* with a mean of 0.23. Among paired differences between ASP and ASPt of our cervical vertebrae sample, the smallest paired difference was 0.01 in both the posterior cervical of *C. frugilegus* and an anterior cervical of the duck *Anas crecca*, indicating an almost complete lack of soft tissues within the internal cavity of these pneumatic elements. As mentioned in the methods, we do not count the neural canal space in the total bone volume of vertebrae, and therefore do not consider the soft tissues and potential air (i.e. paramedullary diverticula; [40]) in the neural canal as part of the internal cavity. This space was fully occupied by the spinal cord across most of our sample (see supplementary Figs. S3 and S4). Notably, the cervical vertebra of *C. frugilegus* is the only one in our dataset to exhibit clear evidence of paramedullary diverticula, with the cervical vertebrae of the ducks *Callonetta leucophrys* and *Anas crecca* exhibiting a small amount of air in the neural canal space that may also correspond to paramedullary diverticula, though this identification is ambiguous (see supplementary material). The greatest paired difference was 0.16 in an anterior cervical of *A. brachyrhynchus*, with the mean of the differences among cervical vertebrae examined being 0.05.

For thoracic vertebrae (*n* = 5 from five different species), ASP ranged from 0.22 in *N. auritus* to 0.45 in *C. frugilegus* with a mean of 0.34. ASPt of the same sample of thoracic vertebrae ranged from 0.16 in the same specimen of *N. auritus* to 0.44 in the same specimen of *C. frugilegus* (Fig. 2). The smallest paired difference between ASP and ASPt of thoracic vertebrae was 0.00 (rounded to two decimal places) in *C. frugilegus*, which, similar to our findings for the cervical vertebrae of this specimen, indicates a lack of detectable soft tissues within the internal cavity. The greatest paired difference was 0.06 in *N. auritus*, with the mean difference among thoracic vertebrae being 0.03. Similarly to the femora investigated, our small sample size precludes a statistical assessment of the relationship between ASP and ASPt among vertebrae. However, there also seems to be a general trend of a direct relationship between ASP and ASPt among both thoracic (zero rank changes) and cervical vertebrae.

### (c) Complete bone versus diaphysis-only comparisons

We compared ASP data between complete long bones versus just the diaphysis (shaft) to determine whether fragmentary long bones may yield comparable estimates to data from complete elements. The greatest paired difference between ASP of the complete humerus and the diaphysis of the humerus was 0.06 in *Corvus frugilegus* (where ASP is greater in the complete humerus than the diaphysis, and therefore implicitly greater in the epiphyses), and in the opposite direction the greatest difference was −0.13 in *Ensifera ensifera* (where ASP is greater in the diaphysis than the complete humerus, and therefore implicitly lower in the epiphyses). A paired samples *t*-test between ASP of the complete humerus and ASP of just the humeral diaphysis (*n* = 38 from 36 different species of bird) showed no significant difference (paired *t*-test: *t*(37) = −0.85831, p = 0.3962), with the mean of the differences being −0.005. This indicates that the proportion of air (i.e. internal cavity in this case, and therefore inversely, the proportion of bone) is generally similar between the humeral diaphysis and the whole humerus in birds. Additionally, this implies that the ratio of bone to internal cavity among just the humeral diaphysis is, on average, almost exactly the same as the ratio of bone to internal cavity of the epiphyses, which is made clear by our findings that for humeri in our dataset, 71% of the total internal cavity by volume and 71% of the total bone by volume, on average, is situated in the epiphyses, with the remaining 29% of both internal cavity and bone volumes residing in the diaphysis.

The same comparisons for the ASPt measure showed the greatest paired difference between the complete humerus and diaphysis of 0.15 in one specimen of the cockatiel *Nymphicus hollandicus* (where ASPt is greater in the complete humerus than the diaphysis, and therefore implicitly greater in the epiphyses). In the other direction the greatest difference was −0.17 in the weaver *Euplectes hordeaceus* (where ASPt is greater in the diaphysis than the complete humerus, and therefore implicitly lower in the epiphyses). A paired samples *t*-test between ASPt of the complete humerus and ASPt of just the humeral diaphysis (*n* = 38 from 36 different species) detected a significant difference (paired *t*-test: *t*(37) = −4.9065, p = 1.883e-05), with the mean of the paired differences being −0.06 (CI 95%: −0.08 to −0.04), and Cohen’s effect size value (*d* = 0.50) indicated a medium effect. These findings show that among pneumatic avian humeri ASPt is greater, on average, in the diaphysis compared to the complete element by about 0.06, which indicates a greater proportion of soft tissues being found in the epiphyses than in the diaphysis. Indeed, this conclusion may be made clearer through comparisons between the distributions of volumes of the components for humeri in our dataset. We find that 67% of the total air by volume, on average, resides in the epiphyses, with the remaining 33% being found in the diaphysis (relatively similar to the distribution expected based on the total internal cavity distribution). However, for soft tissues, 86% of the total soft tissue volume, on average, resides in the epiphyses, with only 15% of the total volume residing in the diaphysis.

We conducted a similar analysis on the diaphysis of femora, allowing us to make pairwise comparisons with the complete femur. The greatest paired difference in ASP between complete femora and just their diaphysis was −0.05 in *Phasianus colchicus*, with the mean difference being −0.03. This negative value indicates a trend towards a slightly higher bone proportion at the epiphyses of pneumatic femora than their diaphysis, though this difference is relatively small. Additionally, 63% of the total bone by volume for femora, on average, was found to reside in the epiphyses, with the remaining 37% of the total bone by volume residing in the diaphysis. In comparison, 58% of the total internal cavity of femora by volume, on average, were found to reside in the epiphyses, with the remaining 42% residing in the diaphysis. For the paired differences in ASPt of complete femora versus just their diaphysis, the greatest difference was −0.09 in one representative of *L. lagopus scotica*. The mean of the differences in ASPt was −0.05, which, similar to our findings for the humerus, indicates a greater proportion of soft tissues being found in the epiphyses of the femora compared with the diaphysis. On average, 56% of the total air by volume for femora were found in the epiphyses, with the remaining 44% residing in the diaphysis (similar to the expected distribution based on the total internal cavity distribution). In comparison, 64% of the total soft tissues by volume for femora were located in the epiphyses, with the remaining 36% residing in the diaphysis.

## Discussion

Previous studies attempting to characterise variation in the extent of pneumaticity generally fall into two main categories. One focus has been on understanding patterns of pneumaticity across the entire skeleton, predominantly by scoring the presence or absence of features indicative of skeletal pneumaticity [e.g., 12, 13, 24, 26, 28, 29, 31]. The other focuses on attempting to estimate the extent of pneumaticity of specific skeletal elements, generally using Air Space Proportion (ASP) as a metric [e.g., 8, 27, 30, 34, 41]. Our study evaluates the extent to which ASP provides a reliable estimate of the true extent of pneumaticity of pneumatic skeletal elements in extant birds, with implications for the reliability of ASP estimates in the fossil record. The importance of using extant specimens with intact soft tissues to validate previous conclusions drawn only from osteological material, and to answer questions related to our holistic understanding of pneumaticity and the pulmonary anatomy associated with this system is highlighted here, and adds to a growing list of studies in recent years to recognise this [e.g., 30, 40, 42, 43, 44].

Values of ASP tending closer to 1 (or 100%) represent elements with larger air spaces within them and proportionally less bone. ASP as a measure of pneumaticity has mostly been applied to sauropods [8, 34, 45–51], though data also exist for pterosaurs [41, 52–54] and extant birds [27] (see Table 3 for a summary of previously published estimates of ASP, and supplementary Table S2 for the complete table). Until recently, all ASP data gathered on birds was estimated only in long bones [8, 41] using previously published data [55–57] of a measure of bone thickness known as *K*, the ratio of the inner diameter to the outer diameter. This can only be applied to tubular bones (i.e. long bones), with *K^2^* providing an estimate of ASP in pneumatic elements. Moore [27] was the first to present ASP values calculated from 3D volumetric data of bird vertebrae; however, that study focused specifically on storks (Ciconiidae), and as such the values from that study may not necessarily be representative of ASP values across the avian crown group.

**Table 3.** Summary of previously published mean ASP values by element for different taxonomic groups. Comprehensive list of all previously published ASP data, including sources, specimen numbers and additional information in supplementary table S2.

| Element | Crown<br>birds | Pterosaurs | Sauropods | Non-avian<br>Theropods |
| --- | --- | --- | --- | --- |
| Caudal vert. |  |  | 0.39 |  |
| Cervical vert. | 0.61 | 0.74 | 0.54 | 0.48 |
| Dorsal/ Thoracic vert. | 0.77 | 0.69 | 0.65 | 0.50 |
| Femur | 0.63 | 0.66 |  |  |
| Humerus | 0.61 | 0.78 |  |  |
| Metacarpal |  | 0.86 |  |  |
| Metatarsal | 0.74 |  |  |  |
| Pelvis |  |  | 0.62 |  |
| Phalanges |  | 0.65 |  |  |
| Presacral vert. (unspecified) |  |  | 0.76 |  |
| Radius |  | 0.65 |  |  |
| Rib |  | 0.77 | 0.55 |  |
| Tibia | 0.67 | 0.72 |  |  |
| Ulna | 0.61 | 0.63 |  |  |

Using ASP as an estimate of the extent of bone pneumaticity within an element relies on the assumption that the volume of soft tissues within the internal cavities of pneumatic bones is negligible (i.e. the entire internal cavity is completely air-filled). In support of this assumption, Wedel [8] references a histological study on domestic chickens (*Gallus gallus domesticus*), in which the pneumatic bones are described as being lined with simple squamous epithelium [58]. However, that study also shows soft tissues between the epithelial lining and bone tissue [58: Fig. 17], an observation that has seemingly gone unmentioned in subsequent literature. The birds investigated in that study were within their first year, so it is possible that some of the residual marrow observed histologically may have been subsequently resorbed as the chickens developed to full maturity, but other studies investigating the architecture of pneumatic elements in mature birds have also noted the presence of at least trace amounts of marrow persisting within the internal cavity of pneumatic bones [59, 60, 61; PMO pers. obs., 62]. Canoville et al. [63] suggest this persistence of marrow from the incomplete invasion of pneumatising diverticula could provide an explanation for the presence of medullary bone in some pneumatic elements. Furthermore, Burton et al. [30] recently documented variable amounts of soft tissue within the internal cavities of pneumatic extant bird humeri, suggesting the possibility that ASP systematically inflates estimates of skeletal pneumaticity, yielding misleading inferences of the extent of skeletal pneumaticity in fossil taxa.

In this study, a comparison of air space proportion (ASP) and true air space proportion (ASPt), accounting for soft tissues within the internal cavity, indicates a significant difference in estimated values. Among the investigated elements, the greatest differences between ASP and ASPt are observed in the humerus (*n* = 38), with an average difference of 0.14 (Table 2). In the femur (*n* = 5) and cervical vertebrae (*n* = 9), estimates of ASP decline on average by a factor of 0.08 and 0.05, respectively, while differences within thoracic vertebrae (*n* = 5) are smaller (0.03). The smaller change in ASP to ASPt among sampled vertebrae may suggest that pneumatic vertebrae genuinely exhibit proportionally less soft tissue within the internal cavities as compared to pneumatic elements of the appendicular skeleton. Unfortunately, however, our limited sample size precludes us from drawing more robust conclusions regarding the extent of skeletal pneumaticity within the avian vertebral column, and we encourage future work investigating differences between ASP and ASPt in this region of the skeleton. This latter point is particularly relevant when considering that skeletal pneumaticity estimates in most groups of extinct archosaurs (e.g., non-avian dinosaurs) are almost exclusively focused on the postcranial axial skeleton [8, 9, 14, 15].

We observe a few instances in which the internal cavity of a pneumatic bone is almost completely air-filled, consistent with the assumptions of ASP [e.g., see 8]. This is true for at least one representative of each of the skeletal elements investigated — the femur of *Phoenicopterus roseus* (ASPi = 100%), the humerus of *Anas crecca* (ASPi = 99%) and an anterior cervical vertebra from the same specimen (C5; ASPi = 97%), a posterior cervical vertebra of *Corvus frugilegus* (C11; ASPi = 98%) and a thoracic vertebra from the same specimen (T3; ASPi = 99%) — indicating that, in some cases, soft tissue content within pneumatic elements may essentially be negligible. However, our results suggest that this is uncommon, with the general trend emerging indicating a significant decrease in the extent of pneumaticity between ASP and ASPt, at least in appendicular elements. This could have important consequences for estimates of bulk skeletal density and body mass of taxa with pneumatic skeletons in the fossil record.

### (a) Implications for mass estimation

Mass estimation in fossil archosaurs exhibiting postcranial skeletal pneumaticity has often been contentious, particularly with regard to sauropods, the largest terrestrial animals of all time [33]. Although various methods for estimating body mass exist, several studies attempting to estimate the mass of sauropods have opted to use volume-based methods [e.g., 8, 64]. Having accounted for ASP of sauropod vertebrae, Wedel [8] suggested that sauropods may have exhibited a whole body bulk density of roughly 0.8 g cm^-3^, and that previous volume-based mass estimates of sauropods should therefore be reduced by approximately 10% due to their extensively pneumatic axial skeletons. However, our findings suggest that this estimated bulk density may be an underestimate based on the assumption of negligible soft tissues within the internal cavities of pneumatic elements. This would be in agreement with Sander et al. [33] who noted that a higher specific density than 0.8 g cm^-3^ would be expected based on allometry of inner ear semicircular canals [65]. Based on our estimates in birds, humeri exhibit average bulk densities of 0.887 g cm^-3^ when soft tissues within internal cavities are unaccounted for, versus 1.025 g cm^-3^ once soft tissues within the internal cavity are considered (Table 2), resulting in a mass increase of approximately 15%. This would also suggest an average estimated mass increase of about 12% among studied femora, 3% among cervical vertebrae and 2% among thoracic vertebrae.

If we take the same logic and apply it to sauropods, in which cervical vertebrae have a mean estimated ASP of 0.54 (Table 3), we might expect, based on our results for cervical vertebrae in this study, that accounting for within-bone soft tissue content would result in an estimated decrease of 0.05 for its equivalent ASPt value, yielding an ASPt estimate of 0.49. Assuming the same densities for bone and marrow as stated in our methods, bulk density of sauropod cervical vertebrae according to ASP would be 0.943 g cm^-3^ and bulk density according to ASPt would be 0.993 g cm^-3^ (Fig. 3). Tentatively, this could indicate an estimated increase in bulk density and mass by about 5% for this element. When accounting for numerous pneumatic elements within a specimen, this difference could meaningfully affect skeletal and total body mass estimation. Here, our example focuses on sauropods, but similar conclusions would be applicable to pterosaurs and any other fossil archosaurs with extensively pneumatised skeletons [66]. Of course, this example comes with its own caveats such as assuming the bone and marrow density in sauropods is similar to that of birds, and assuming that the proportion of soft tissues within the internal cavities of pneumatic cervical vertebrae would scale isometrically between birds and sauropods, despite sauropod vertebrae being several orders of magnitude larger. Nevertheless, these findings demonstrate the extent to which the systematic overestimation of skeletal pneumaticity from ASP may impact estimates of bulk density, and ultimately both skeletal and total body mass, emphasising the need for further investigation of this topic.

**Figure 3.**
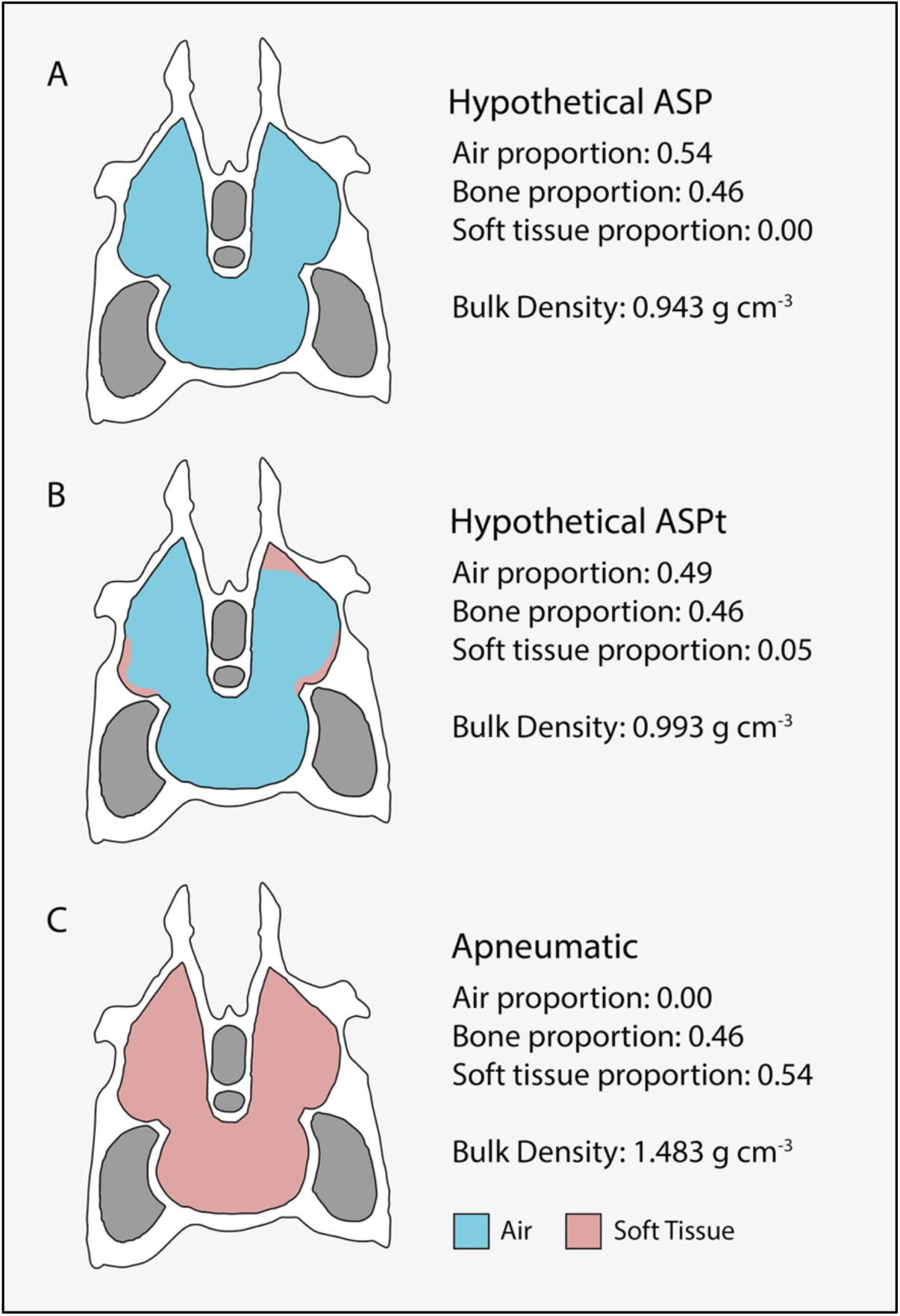
Hypothetical scenarios of varying degrees of pneumaticity in sauropod vertebrae and their implications on bulk density estimates. (A) Pneumatic sauropod vertebra with no soft tissues in the internal cavity (as assumed in the air space proportion measure, ASP). (B) Pneumatic sauropod vertebra with hypothetical proportion of soft tissues in the internal cavity based on our estimates for extant birds (i.e. a hypothetical true air space proportion, ASPt). (C) Sauropod vertebra if the internal cavity were completely filled with soft tissues, included here as an illustrative example. *Diplodocus* vertebra diagram modified from Taylor and Wedel [34], figure 4.3 and not to scale. Colours represent bone (white), air (blue), soft tissues (pink) and non-intraosseous areas (grey). ASP value in (A) represents mean sauropod cervical vertebra ASP as calculated from previously published estimates (see table 3 and supplementary table S2).

### (b) Continued importance of ASP data

Despite our findings that ASP estimates based on the original framework of assuming the absence of soft tissues may not be representative of ASPt, ASP data remain relevant and provide important insights into the extent of an organism’s skeletal pneumaticity. By definition, ASP represents the upper bound of an element’s pneumatisation. Our Spearman rank results also show that ASP strongly correlates with ASPt (ρ(36) = 0.758, p = 3.314e-07), indicating that for a specific element, the relative degree of pneumaticity of individual elements is likely to be similar for both ASP and ASPt.

There is evidence to suggest that pneumatising diverticula invade bone in an opportunistic manner and are essentially modulated by constraints imposed by local biomechanical loading regimes [27, 67]. The increased number of bony trabeculae at the epiphyses of long bones may help restrict the penetration of pneumatic diverticula, resulting in an increased likelihood of some pockets of marrow remaining among spaces between trabeculae where pneumatic diverticula were unable to invade. If this is true, it might also suggest that pneumatic bones with a greater proportion of trabeculae might also have a greater proportion of soft tissues, though additional data using intact and fresh avian specimens will be necessary to evaluate this hypothesis. Additionally, this idea may also be important in the context of associating variable amounts of soft tissue between vertebrae classified as camerate (with the internal structure made up of few and large internal chambers) and camellate (with the internal structure made up of more numerous and smaller internal chambers separated by thin bone; [6]), whereby vertebrae with more camellate morphologies may have contained a greater fraction of soft tissues.

ASP may still be a powerful indicator of the functional aspects of a pneumatic bone and what species with pneumatic skeletons may have been functionally capable of, especially as it allows us to make consistent comparisons among both extant taxa and specimens from the fossil record. Indeed, estimating maximum ASP for a given element may be our only option for interpreting the extent of pneumaticity in a fossil element, where soft tissues are not preserved. Osteological correlates of pneumaticity are indicators of where soft tissues from the pneumatic system (e.g., air sacs and diverticula) have interacted with and left marks on the skeleton. Identification of these markers is important for inferring the presence of pneumaticity in fossil elements; however, these osteological correlates do not provide a quantitative indication of the extent of pneumaticity of a specific element. Such osteological correlates include pneumatic foramina (relatively large skeletal perforations associated with the invasion of pneumatic diverticula), which are the most widely accepted unambiguous osteological correlate of PSP [10, 14], as well as ‘pneumosteum’ (fibrous markers associated with pneumatic diverticula attachment sites on secondary endosteal trabecular bone; [20]). Because the latter correlate is associated with pneumaticity within the internal cavity of a bone, identification of areas where pneumosteum is present or absent within the same element could potentially be indicative of the distribution of soft tissues within the internal cavity of the element, as areas lacking pneumosteum could have soft tissues blocking the attachment of pneumatic diverticula to the trabecular bone. However, future work would be necessary to evaluate the utility and accuracy of this approach. Pneumatic fossae, bony laminae and differential surface texture have also been associated with the presence of pneumaticity [8, 10], and may be indicative of an avianlike respiratory system exhibiting air sacs and diverticula; however, the presence of these correlates alone is not indicative of the extent of postcranial skeletal pneumaticity [68].

Furthermore, due to a lack of intraspecific data into the variability of soft tissues within internal pneumatic cavities (few studies have investigated this topic and only among domesticated species; e.g., [59]), it remains unclear whether the extent to which the internal cavities of pneumatic elements become air-filled (i.e. ASPi) is consistent within species. The reliability of accurately estimating ASPt from the fossil record may hinge on this, as ASPi influences the discrepancy between ASP and ASPt. Therefore, investigating this across a greater range of taxa, with multiple representatives of particular species, will clarify the extent to which we can make general assumptions about ASPi (and therefore expected ASP vs ASPt discrepancy).

At present, ASP data remain important and are worth collecting in the context of investigating pneumaticity and its role in bulk density and mass estimation. However, future studies should note that these data represent the extreme upper bound of the extent to which a pneumatic element was air-filled in life, and that pneumatic elements with negligible soft tissue content appear to be uncommon [30; this study].

Our survey of previously published ASP data (Table 3 and supplementary Table S2), revealed that postcranial ASP data on non-avian theropods is all but lacking. Wedel [8] included some ASP estimates for theropod vertebrae based on previously published cross-sections, while others have reported data for cranial elements. For example, Funston et al. (2018) published an ASP estimate for the pneumatic squamosal of a tyrannosaurid [69], though the material was initially incorrectly identified as part of a pterosaur pelvis [70]. Gathering ASP data on non-avian theropods will therefore represent an intriguing area for future research.

### (c) Insights into the skeletal architecture of long bones from ASP and ASPt estimates

Despite the number of assumptions inherent to estimates of ASP in avian long bones derived from the variable *K*, the mean estimate of ASP based on such data reported by Wedel [8] (0.59-0.64) is remarkably accurate with respect to our 3D volume-derived estimates of ASP for the humerus (mean = 0.57) and femur (mean = 0.65). This may be explained by insights into the structure of long bones from our results. Among humeri, a paired *t*-test between ASP of complete long bones and just their diaphyses failed to reach significance. This implies that ASP, and inversely bone proportion, remain virtually the same (0.57; Table 2) between the diaphysis and collective epiphyses of pneumatic humeri in birds, despite these areas exhibiting pronounced structural differences. The diaphysis (shaft) of long bones tends to be uniformly long and tubular with little trabecular bone, while the epiphyses tend to be comparatively wide with thinner cortical bone [e.g., 41], and are often more complex in shape with a web of internal trabecular bone. The consistency in ASP estimates between the diaphysis and epiphyses in avian humeri may therefore indicate that bone proportion lost through a combination of increased internal volume from widening at the epiphyses and thinning of cortical bone may be replaced by an approximately equivalent proportion of trabecular bone. This may be related to biomechanical factors beyond the scope of this investigation, but this hypothesis may represent a fruitful avenue of future research. These results are different from findings on ASP variation in pterosaur wing bones, where ASP has been found to be higher in the epiphyses versus the diaphysis [41]. Among avian femora, the mean difference in ASP between the complete bone and just the diaphysis was −0.03 (Table 2), which, though small, is more pronounced than the differences we observe in avian humeri. This slight negative value indicates a trend towards decreased ASP within the epiphyses compared to the diaphysis in femora. Although more data may be necessary to confirm these findings, our results would suggest that estimating ASP from just the diaphysis of pneumatic humeri may generally be representative of ASP for the entire element, at least among crown birds and their close stem group relatives. This is a relevant and important factor to assess as long bones preserved in the fossil record are often incomplete, taphonomically deformed and matrix-filled, which can be particularly problematic at their epiphyses (e.g., the deltopectoral crest of fossil humeri are commonly fragmented or missing; [71]). Furthermore, the delicate structure of trabecular bone at epiphyses is difficult to segment in CT scans of fresh extant bird specimens, and potentially even more challenging in fossil specimens where matrix can infill these internal cavities. Therefore, assessing the discrepancies in ASP between complete long bones versus the shaft alone allows us to determine the viability of disregarding epiphyses to facilitate and expand data collection for future estimates of ASP derived from 3D volumetric data. On the other hand, ASPt differs significantly when data for complete humeri are compared to their diaphyses, with the mean differences for humeri being −0.06, and a similar mean difference of −0.05 being found for femora (Table 2). This indicates that a greater proportion of soft tissues are distributed in the epiphyses than in the diaphysis among avian long bones, which has been noted from previous observations [30, 61].

### (d) Limitations of our methodological approach

Validity of the data and results from this study rely on the assumption that the intraosseous material of intermediate density between bone and air observed in CT scans of intact bird carcasses are attributable to soft tissues present in life. However, the nature of these tissues has yet to be validated histologically. The supplemental information of Burton et al. [30] discusses potential sources of false negative and false positive inferences of pneumatisation from CT scans of frozen specimens. These include soft tissue shrinkage from freezing potentially leading to underestimates of soft tissue volume, and the presence of freeze-thaw or decay-related fluids (or ice; [72]) within bones, which have similar densities to soft tissue, potentially leading to overestimates of soft tissue volume. It is of particular importance to highlight the latter point, as recent studies have shown that decay-related accumulation of water-density substances within the lung and pneumatic spaces in birds can occur rapidly after death (within 8 hours; [43, 73 (this volume)]). Thus, time between death and freezing is important to consider when studying the pulmonary system in birds, with one of the most recent investigations of this topic advocating for studies of PSP that assess soft tissues associated with the pulmonary/pneumatic system to only sample specimens that are either live or frozen within 1-2 hours of death [73 (this volume)]. Our study was designed to utilise pre-existing CT datasets (i.e. utilising many of the scans investigated in [30]) of a broad phylogenetic range of specimens that had been salvaged, thus time between death and freezing is unknown among our specimens, although as described by Burton et al. [30], we attempted to exclude specimens exhibiting clear signs of decay or freezerburn based on visual assessment of CT scans. This may have important consequences for our estimates of ASPt, particularly when considering that decay-related fluids cannot be readily distinguished from genuine intraosseous soft tissues in CT scans as they both fall within the same water-density grey-value range. Considering the results presented by Moore and Schachner [73 (this volume)], it seems that several specimens in our dataset may indeed present intraosseous material that is consistent with decay-related fluid (see supplementary figures S1-5), which has likely resulted in inflated volumes of ‘marrow’. The effects of this on ASPt would be a smaller estimated value, and a more dramatic ASP to ASPt paired difference. Although the present study provides preliminary results and an experimental framework for future studies to continue assessing variations in marrow volume that may persist in pneumatic elements, we acknowledge that our results will ideally need to be validated using CT scans of fresh specimens in which the time between death and freezing has been carefully controlled for (ideally within 1-2 hours).

It is also noteworthy to mention that other studies that have utilised birds with soft tissues intact to investigate the pulmonary/pneumatic system have artificially inflated the specimens prior to scanning [e.g., 42–44, 73 (this volume)], which was not a methodological step taken here. This step is necessary when studying lungs, air sacs and pneumatic diverticula [43]. However, the effects of artificial inflation on pneumatising diverticula, as compared to simply scanning frozen specimens without artificially inflating them, have not been explicitly investigated, thus it is unknown how this may affect estimates of ASPt. Future studies may seek to validate the method used here by making direct comparisons of ASPt estimates on freshly frozen specimens when scanned with and without artificial inflation.

Additionally, some studies opt to use contrast staining methods (e.g., iodine) prior to CT scanning in order to enhance contrast and differentiate soft tissues in CT scan data [72]. However, the use of contrast staining methods were not considered here as it would be time-consuming and costly to stain ∼40 adult birds, which also require fixation prior to scanning [72]. Furthermore, the process of fixation and staining can also cause soft-tissue shrinkage [72, 74], is more destructive to the specimen than simply freezing, and also would result in fluid infilling pneumatic spaces, which would itself result in unreliable ASPt data. Thus, contrast-enhanced staining methods in this case could potentially introduce more uncontrollable factors than simply CT scanning intact frozen specimens. Future studies may want to test the validity of the method used here by performing methodological comparisons using the same specimen. For instance, CT scanning a whole, intact and fresh specimen before and after freezing, and then collecting histological samples or performing contrast-enhanced CT scanning methods to confirm that the presence of intraosseous soft tissues aligns with inferences from scans. The non-destructive techniques used in this study are justified on the ability to perform broad-scale comparisons using many specimens in a timely manner, while preserving them intact for future studies.

## Conclusions

Collecting data on the air space proportion (ASP) measure following its original definition continues to be useful and relevant to our understanding of the evolution of skeletal pneumaticity. This is particularly important for palaeontologists, as it provides a means to compare upper limits of the extent to which a pneumatic element was air-filled in life. However, it is important to recognise that the extent to which the internal cavity of a given element may have been air-filled would not have been 100%, with the true air space proportion (ASPt) likely to have been significantly lower. With data collection on the variation of soft tissues within the internal cavity of pneumatic bones at only an incipient stage, we cannot predict the ASPt of fossilised pneumatic elements with precision, and data generated to date have illustrated considerable interspecific variation. Moreover, intraspecific variability remains essentially uninvestigated. Nonetheless, our data allow us to make generalised predictions about the expected decrease in air space proportion between ASP and ASPt, providing a preliminary basis for constraining such estimates in fossils.

We also find that ASP estimated from just the diaphysis of a pneumatic long bone generally provides an accurate representation of ASP for the entire element. This may be useful when attempting to estimate ASP from the fossil record based on partial elements, or where it is difficult to distinguish and segment trabecular bone in the epiphyses due to matrix infill. With more structurally complex elements (e.g., vertebrae), it may still be best practice to estimate ASP from complete 3D volumetric measurements when possible, or calculate an average from multiple 2D slices [e.g., 47].

The data presented herein adds to the growing sample of pneumaticity data in the form of ASP estimates and represents the first based on 3D volumetric data from avian long bones. Among other fruitful directions, we suggest that future studies should aim to broaden our understanding of intraspecific variation in ASPt among extant birds to help clarify the extent to which skeletal pneumatisation may have been adaptively associated with mass reduction or skeletal volume increases in extinct archosaurs.

## Acknowledgements

This work was funded by UKRI grant MR/X015130/1 to DJF. This work was also supported by the University of Cambridge Harding Distinguished Postgraduate Scholars Programme and by the Natural Environment Research Council grant number NE/S007164/1 to MGB. For the purpose of open access, the authors have applied a Creative Commons Attribution (CC BY) licence to any Author Accepted Manuscript version arising.

